# Error-suppression mechanism of PCR by blocker strands

**DOI:** 10.1101/2022.06.02.494611

**Authors:** Hiroyuki Aoyanagi, Simone Pigolotti, Shinji Ono, Shoichi Toyabe

## Abstract

The polymerase chain reaction (PCR) is a central technique in biotechnology. Its ability to amplify a specific target region of a DNA sequence has led to prominent applications, including virus tests, DNA sequencing, genotyping, and genome cloning. These applications rely on the specificity of the primer hybridization, and therefore require effective suppression of hybridization errors. A simple and effective method to achieve that is to add blocker strands, also called as clamp, to the PCR mixture. These strands bind to the unwanted target sequence, thereby blocking the primer mishybridization. Because of its simplicity, this method is applicable to a broad nucleic-acid-based biotechnology. However, the precise mechanism by which blocker strands suppress PCR error remains to be understood, limiting the applicability of this technique. Here, we combine experiments and theoretical modeling to reveal this mechanism. We find that the blocker strands both energetically destabilize the mishybridized complex and sculpt a kinetic barrier to suppress mishybridization. This combination of energetic and kinetic biasing extends the viable range of annealing temperatures, which reduces design constraint of the primer sequence and extends the applicability of PCR.

## Introduction

PCR is used in a broad, ever-expanding range of biotechnological applications^1^. Fidelity of PCR is determined by the specificity of the primer hybridization. In applications, mishybridization leads to unwanted consequences, such as false positives in virus tests and sequencing errors. Given the importance and widespread nature of these applications, methods for suppressing hybridization errors are crucial. A simple and effective method to suppress mishybridization is to add blocker strands, also called as clamp, to the reaction mixture^2–5^. The blockers bind to the target mutants and suppress mishybridization to the mutant sequence. The hybridization of nucleic acids is essential in many biotechnology such as genome editing^6^, DNA nanotechnologies^7^, and RNA interference^8^. The blocker method may be applicable to these systems.

However, despite its effectiveness, the detailed mechanism by which blockers suppress errors remains unclear. This limits the rational design of the blocker sequence^4^. One hypothesis is that the blocker strands energetically destabilize the mishybridized state. An alternative hypothesis is that the blocker strands effectively create a kinetic barrier to kinetically suppress the mishybridization. Discerning between these hypotheses would improve our understanding of how PCR works, and suggest design principles for optimized protocols. In this paper, we combine experiments and theory to show that PCR blockers suppress errors in PCR by a combination of energetic and kinetic discrimination. A non-trivial consequence of our theory is that, in the presence of blockers, the range of viable annealing temperatures for the PCR reactions is considerably broadened.

To illustrate the factors that determine the hybridization error, we consider the example of a PCR mixture containing a right template R and a contaminated wrong template W with a mutation in the primer binding region (Fig. 1a). During a PCR cycle, the temperature is lowered from a high denaturing temperature to the annealing temperature *T*_a_. Then, a primer strand P hybridizes to either R or W. Since the primer hybridization is reversible, P repeatedly hybridizes to or dissociates from the template. Eventually, a polymerase binds to the hybridized complex P: R or P: W and elongates P to produce a complementary copy 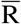 or 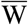, respectively. Important quantities characterizing the PCR are the growth rates *α*_R_ and *α*_W_ 1 and error rate *η*, defined by

**Figure 1.**
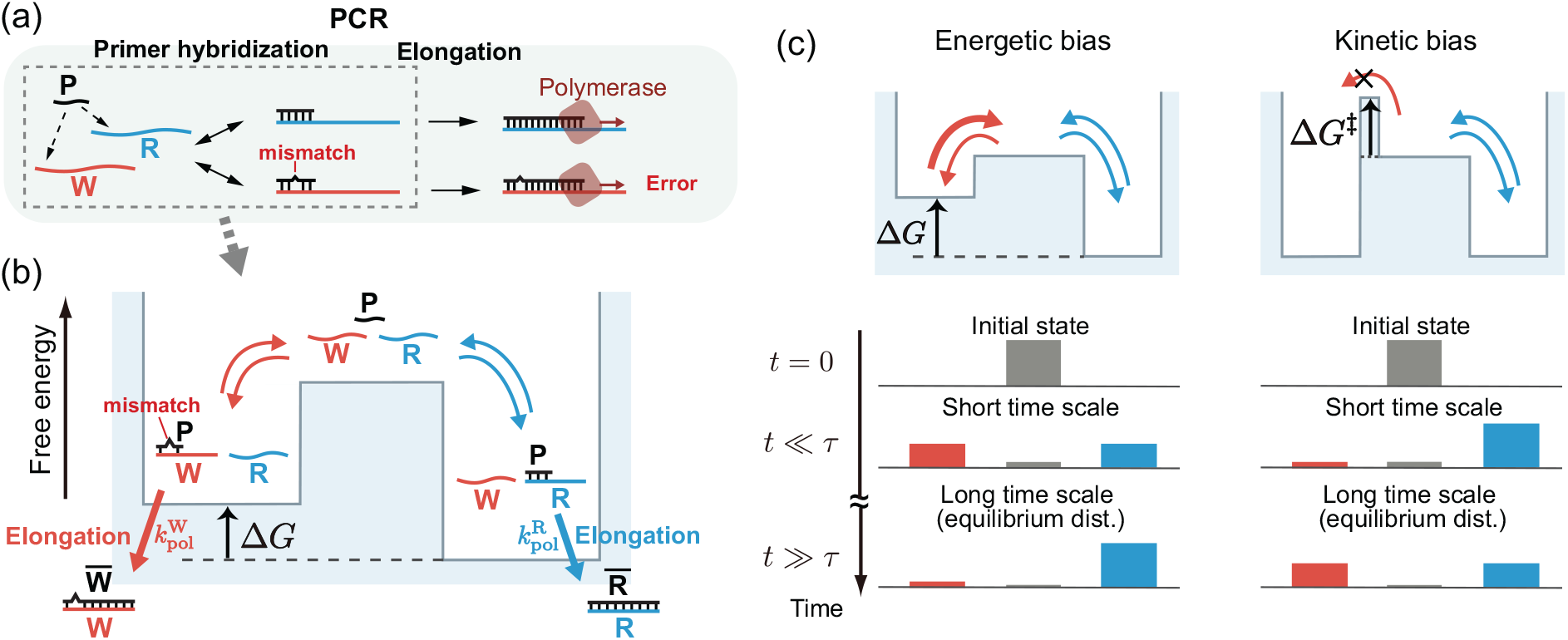
Energetic versus kinetic biasing in PCR. (a) Standard PCR scheme. P, R, and W are the primer, right template, and wrong template with a mismatch in the primer-binding region, respectively. (b) Energy landscape corresponding to the primer hybridization in PCR. Here, 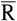 and 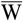 denote the right and wrong products, respectively. (c) Energy landscape for energetic (left) versus kinetic (right) biasing. At an early time, [P: R] and [P: W] are similar because of the similar barrier height for the hybridization. In the bottom, the colored bars correspond to the fractions of [P: R] (orange), [P: W] (blue), and the free templates (gray). At thermodynamic equilibrium, the amount ratio of the wrong hybridization to the right one is 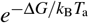, where ∆*G* is the energetic bias and *T*_a_ is the annealing temperature. In the figure, ∆*G*^‡^ is the kinetic bias and *τ* is the relaxation time of the binding dynamics.

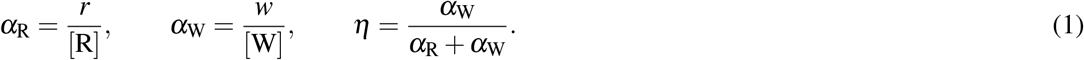

Here, [·] denotes a concentration and *r* and *w* are the increases in concentrations of 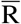 and 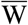 in a cycle, respectively. Hence, *α*_R_ and *α*_W_ represent the fractions of copied strands per template in a cycle. Ideally, one wants to maximize the growth rate *α*_R_, also called the PCR efficiency, and at the same time minimize the error rate *η*.

The accuracy of conventional PCR relies on primer hybridization to R, being energetically more stable than hybridization to W, as quantified by the free energy difference ∆*G* between P: R and P: W (Fig. 1b). This energetic bias can be increased to reduce *η* by carefully designing the primer sequence and increasing the annealing temperature *T*_a_^1^.

However, this approach has an inherent limitation. To see that, we consider the hybridization kinetics (Fig. 1b). The DNA binding rate is usually diffusion-limited and thus does not significantly depend on the sequence^9,10^. Hence, we assume that P hybridizes to R and W at the same rate. On the other hand, the dissociation rates depend on ∆*G*. Repeated hybridization and dissociation of P eventually bring the system to thermodynamic equilibrium, where the amount ratio of the wrong hybridization to the right one is 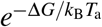. Here, *k*_B_ is the Boltzmann constant. Thus, the error rate in this equilibrium limit is equal to (SI Section S4). 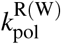 are the elongation rates on R and W by the polymerase. In this case, one can show that the short time error rate is always larger than *η*_eq_^11^, see Fig. 1c. In fact, one problem with this approach is that the enzymatic reaction is usually quite efficient and starts elongation before the binding equilibrates. This means that the error rate is usually not as small as one would expect from Eq. (2). Slowing down of the reaction by, for example, reducing polymerase concentration would lower the error rate by allowing sufficient time for equilibration, but at the cost of efficiency.

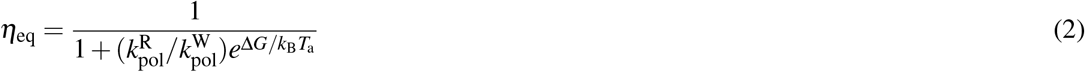

An alternative strategy is to sculpt a kinetic bias by building asymmetric barriers characterized by a difference ∆_*G*_^‡^, so that P preferably binds to R (Fig. 1c, right). Theory predicts that such kinetic bias can reduce *η* without sacrificing efficiency^11^.

### Brief methods

To quantify the error rate, we perform PCR with only a single side of the primer set (Fig. S1). Hence, the product concentration increases linearly, rather than exponentially as in standard PCR (Fig. S2). We mix a primer strand P, two variants of 72-nt template DNA strands R and W, indicated concentrations of thermostable DNA polymerase, and necessary chemicals for the reactions. We also add blocker strands depending on the experiments. The R and W templates are mixed at the same concentrations ([R] = [W] = 2.5 nM), much smaller than the primer concentration ([P] = 100 nM). The P strand binds to R without mismatches and to W with a single-base mismatch. In these conditions, the hybridization error is expected to be large. After hybridization, polymerases copy the template and produces 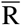 or 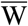. We repeat 10 or 40 thermal cycles to reduce statistical errors, measure *r* and *w*, and calculate the error rate and the efficiency by means of Eq. (1).

The blocker strands B_R_ and B_W_ are 16-nt chimeric strands of DNA and locked nucleic acids (LNA) bases. They hybridize to the primer-binding region of R and W. The B_R(W)_ strand hybridizes to R(W) without mismatches and to W(R) with a single mismatch. Two bases at the 3’ end of the blocker strands are floating to prevent them from acting as primers. Blocker hybridization to the template is faster and more stable than primer binding to the template because of their high concentration ([B_R(W)_] = 20[P] = 2000 nM) and the four LNA bases placed in the vicinity of the mismatch position, which significantly increases hybridization specificity^12,13^.

## Results

### PCR in the absence of blocker strands

We first characterized the performance of conventional PCR by measuring the efficiency *α*_R_ and error rate *η* as a function of the annealing temperature *T*_a_. In the absence of the blocker strands, *α*_R_ and *α*_W_ were large at low *T*_a_ (Fig. 2a, b). As *T*_a_ increased, *α*_R_ decreased at *T*_a_ exceeding the melting temperature of 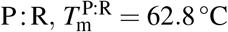. *T*_m_ is defined as the temperature where half of the DNA strands form the complex. On the other hand, *α*_W_ decreased at 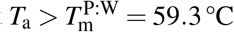. Accordingly, the error rate *η* was large at low *T*_a_ and decreased when 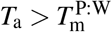 (Fig. 2c). Our results confirm that, in conventional PCR, *T*_a_ needs to be finely tuned in the range 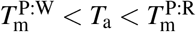 for simultaneously achieving high *α*_R_ and low *η*.

**Figure 2.**
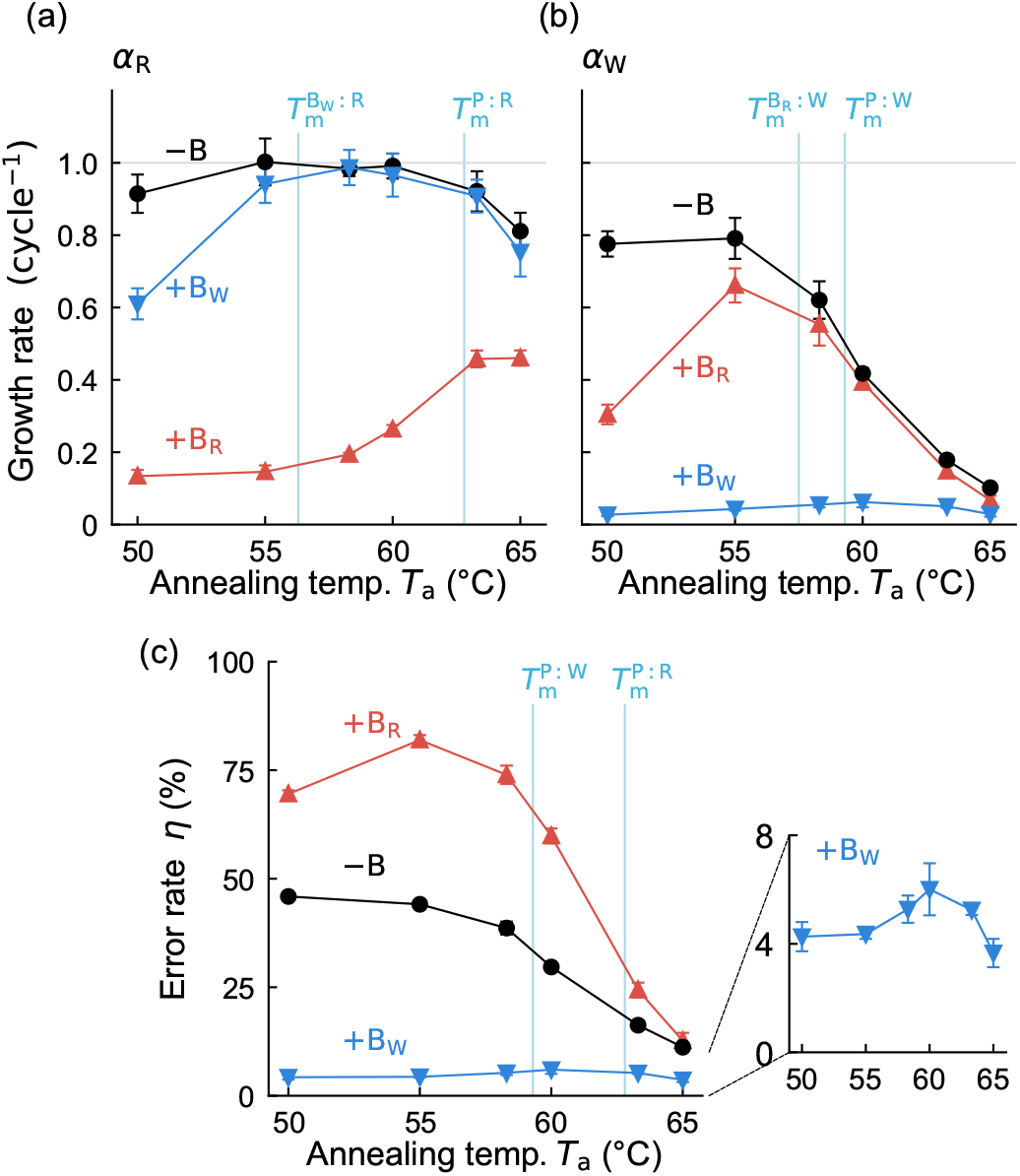
PCR efficiency and error as a function of the annealing temperature. The efficiencies of producing 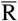 (a) and 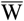 (b) as the function of the annealing temperature *T*_a_. (c) Error rate *η* calculated by Eq. (1). The inset is the magnification of +B_W_. The error bars indicate the standard deviations. Polymerase concentration is 25 units*/*mL, as in the standard PCR protocol.

### Error suppression by blocker strands

We next studied the effect of the blocker strands on the efficiency and the error rate. Intuitively, we expect the blockers to affect the PCR dynamics in the following way. The blocker B_W_ preferably hybridizes to W. As the temperature is lowered to *T*_a_ during the thermal cycle, B_W_ should quickly occupy most W while binding to a small fraction of R only. Hence, the hybridization of P should be significantly biased towards R, thus suppressing the error without sacrificing the speed. On the other hand, the addition of B_R_ prevents P from hybridizing to R. Therefore, we expect an increased error rate in this case.

Indeed, the addition of B_W_ drastically suppressed the errors at all the annealing temperatures we tested (Fig. 2c) without significantly impairing efficiency, at least for large *T*_a_ (Fig. 2a). At 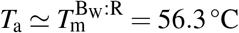, *α*_R_ was reduced due to the hybridization of B_W_ to R. We found that the errors are still effectively suppressed at *T*_a_ much lower than 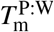, meaning that fine tuning of *T*_a_ is not needed in the presence of blockers.

In contrast, B_R_ drastically reduced *α*_R_ and increased *η*. At *T*_a_ *<* 60 °C, *η* was larger than 50 %, meaning that B_R_ inverted the preference of P hybridization. This setup could be used to amplify rare sequences that would be otherwise difficult to sample.

We used chimeric DNA strands containing LNA bases for the blocker strands, which enhance the specificity of hybridization. The blocker strands with only DNA bases had a limited effect when used with the same concentration and strand length as the chimeric strands (Fig. S7).

### Mathematical model

We quantified the PCR kinetics and in particular the role of blockers using a mathematical model (see SI section S4 for details). The model includes reversible hybridization rate of P to the template strands. In contrast, we assume that the blockers are always at chemical equilibrium as their high concentrations make their hybridization and dissociation dynamics very fast. For simplicity, polymerization is modeled as a single rate without explicitly including polymerase binding and dissociation.

Introducing blockers creates an effective kinetic bias, and at the same time enhances the effective energetic bias between right and wrong targets (Fig. 3a). These effects are quantified by

**Figure 3.**
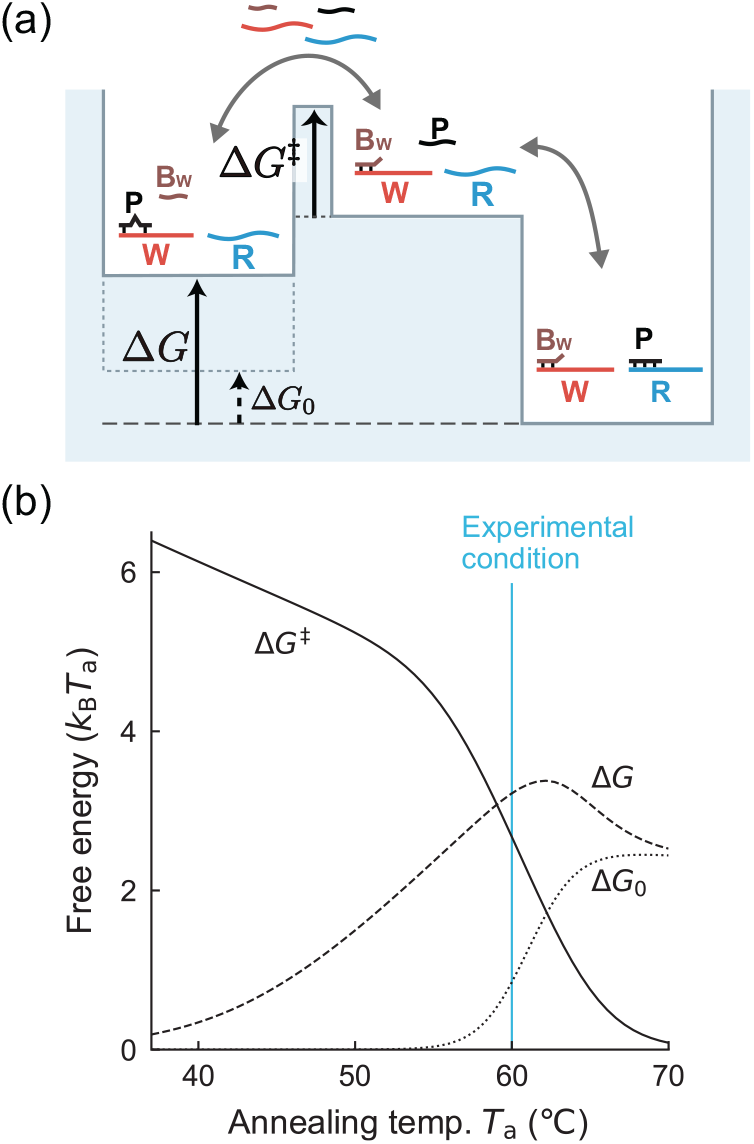
Effective energetic and kinetic bias in the presence of blockers. (a) The free energy landscape in the presence of B_W_. Dependence of energetic (∆*G*, ∆*G*_0_) and kinetic (∆*G*^‡^) bias on *T*_a_ calculated by the model (SI section S4).

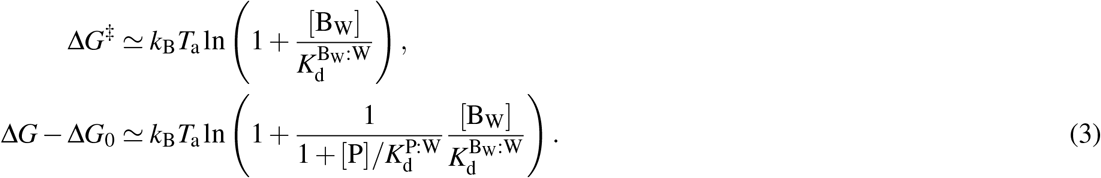

Here, ∆*G*_0_ is the energetic bias in the absence of the blocker, and *K*_d_ is the dissociation constant of the specified hybridization.

We measured the hybridization melting curves and estimated the temperature dependence of *K*_d_ (Fig. S4) to evaluate the dependence of ∆*G* and ∆*G*^‡^ on *T*_a_ (Fig. 3b). We found that ∆*G ≫* ∆*G*^‡^ at high *T*_a_ and ∆*G*^‡^ *≫* ∆*G* at low *T*_a_. This result shows that the error suppression by blocker in a broad *T*_a_ range stems from the combined effects of the energetic and kinetic biasing pronounced at high and low *T*_a_, respectively.

### Kinetics of error suppression

To analyze the detailed kinetics of error suppression by the blocker strands, we varied the polymerase concentration by more than two orders of magnitude while fixing *T*_a_ to 60 °C, which is an appropriate temperature for our primer sequence (Fig. 4). Since elongation by polymerase quenches the hybridization dynamics, a change in the polymerase concentration tunes the time scale available for the hybridization dynamics to relax.

**Figure 4.**
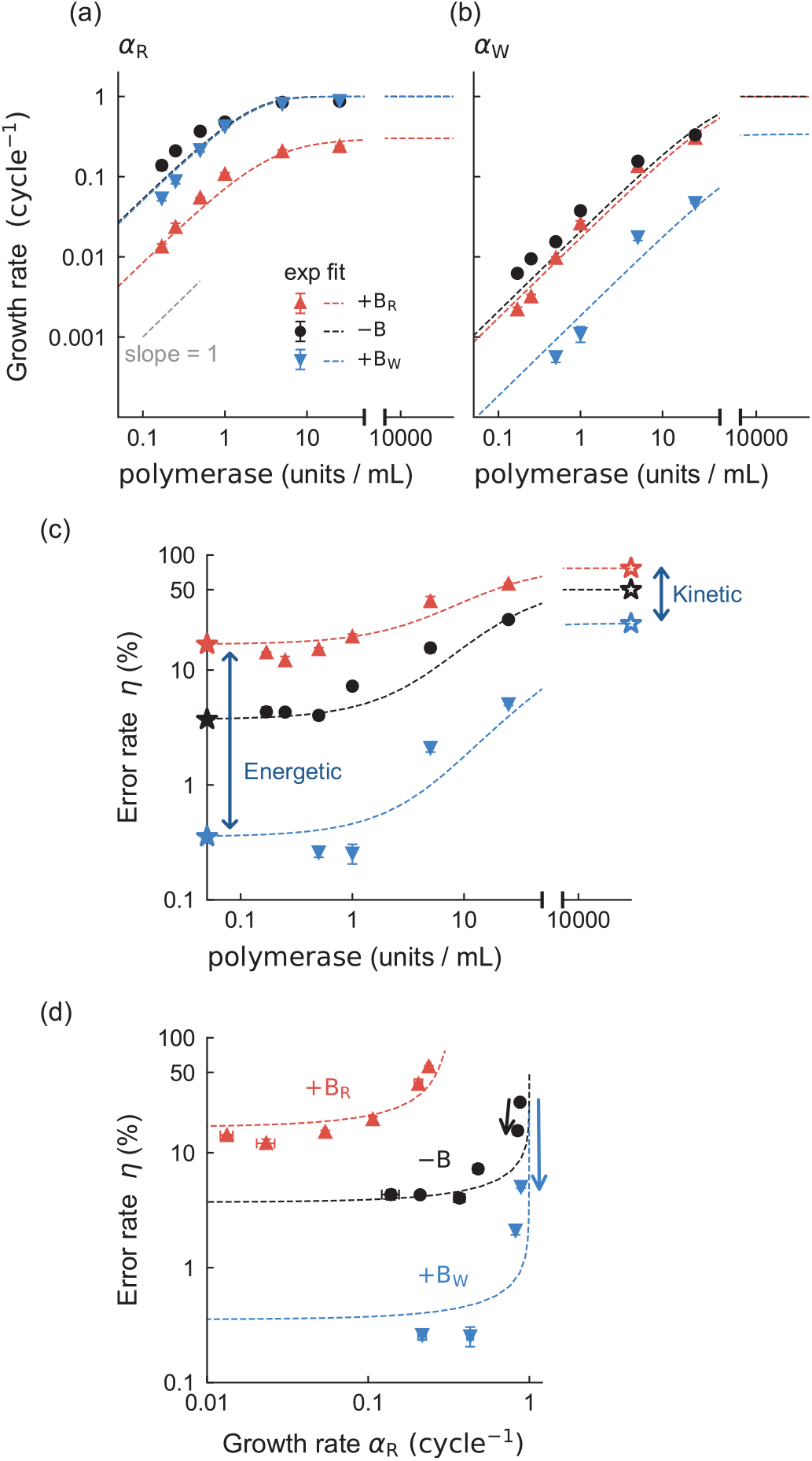
Improved performance of PCR reaction in the presence of blockers is consistent with model predictions. Dependence of the efficiencies (a, b) and error rate (c) on the polymerase concentration. (d) The error rate is plotted against the growth rate *α*_R_ (efficiency). Symbols denote the experimental data, and dashed lines correspond to the model fitting. Stars indicate the limiting values at high (open) or low (closed) polymerase concentrations, which correspond to *η*_eq_ given by (2) and *η*_fast_ given by (4), respectively. The polymerase concentration of the standard PCR protocol is 25 units*/*mL. *T*_a_ = 60 °C. We excluded two points with negative averages due to the statistical errors from (b) and (c) (+B_W_ with 0.17 and 0.25 units*/*mL polymerase). The error bars indicate the standard deviations. See Fig. S5 for the plots in the linear scale.

In the absence of blocker strands, *η* decreased as the polymerase concentration decreased (Fig. 4c). In the low polymerase concentration limit, the limiting error rate *η*_eq_, see Eq. (2), was lower in the presence of blockers B_W_. Since the error is determined by the energetic bias in this limit, this result confirms our prediction that the blocker addition increases ∆*G*.

The blocker addition lowered *η* at high polymerase concentration as well (Fig. 4c), consistently with our prediction that the blockers also create an effective kinetic barrier for the wrong primer strands. The theory approximates the limiting error rate as

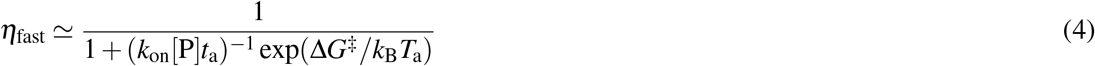

(SI Section S4), depending on the barrier difference ∆*G*^‡^. Here, *k*_on_ is the binding rate of the primer, and *t*_a_ = 30 s is the duration of the annealing step. These results support that the blocker strands suppress errors both energetically and kinetically, as illustrated in Fig. 3a.

The model successfully reproduced the experimental results (dashed lines in Fig. 4). The values of the five fitting parameters were comparable with estimates based on previous work (see SI section S5). Even without blocker strands, a slight reduction of polymerase concentration is effective at suppressing errors without affecting much the efficiency (black arrow in Fig. 4d). However, this strategy requires fine-tuning of the polymerase concentration to maintain the efficiency. On the other hand, the addition of blocker strands is more effective at reducing errors and without reducing the efficiency (blue arrow in Fig. 4d).

### Kinetic regime

Our theoretical model suggests the presence of a novel error correction regime. This fact is related with the relative magnitude of ∆*G* and ∆*G*^‡^, which changes at a temperature *T*_*a*_ *≈* 58 °C where the ∆*G* and ∆*G*^‡^ curves intersect each other in Fig. 3b. According to a theory^11^, the relative magnitude of these two quantities determines the behavior of the primer-binding dynamics.

To illustrate this theoretical prediction, we ran a simulation of our model but without the elongation by polymerase. On the one hand, in the “energetic regime” characterized by ∆*G >* ∆*G*^‡^ (60 °C), the fraction *γ ≡* [P: W]*/*([P: R] + [P: W]) decreases with the annealing time *t*_a_ and converges to an equilibrium value 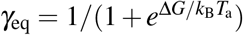 (Fig. 5a). On the other hand, in the “kinetic regime” characterized by ∆*G <* ∆*G*^‡^ (57.5 °C), the kinetic barrier blocks the primer binding to the wrong template (Fig. 1c), and *γ < γ*_eq_ is observed. At the limit of vanishing *t*_a_, *γ* converges to 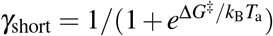 (SI Section S4).

**Figure 5.**
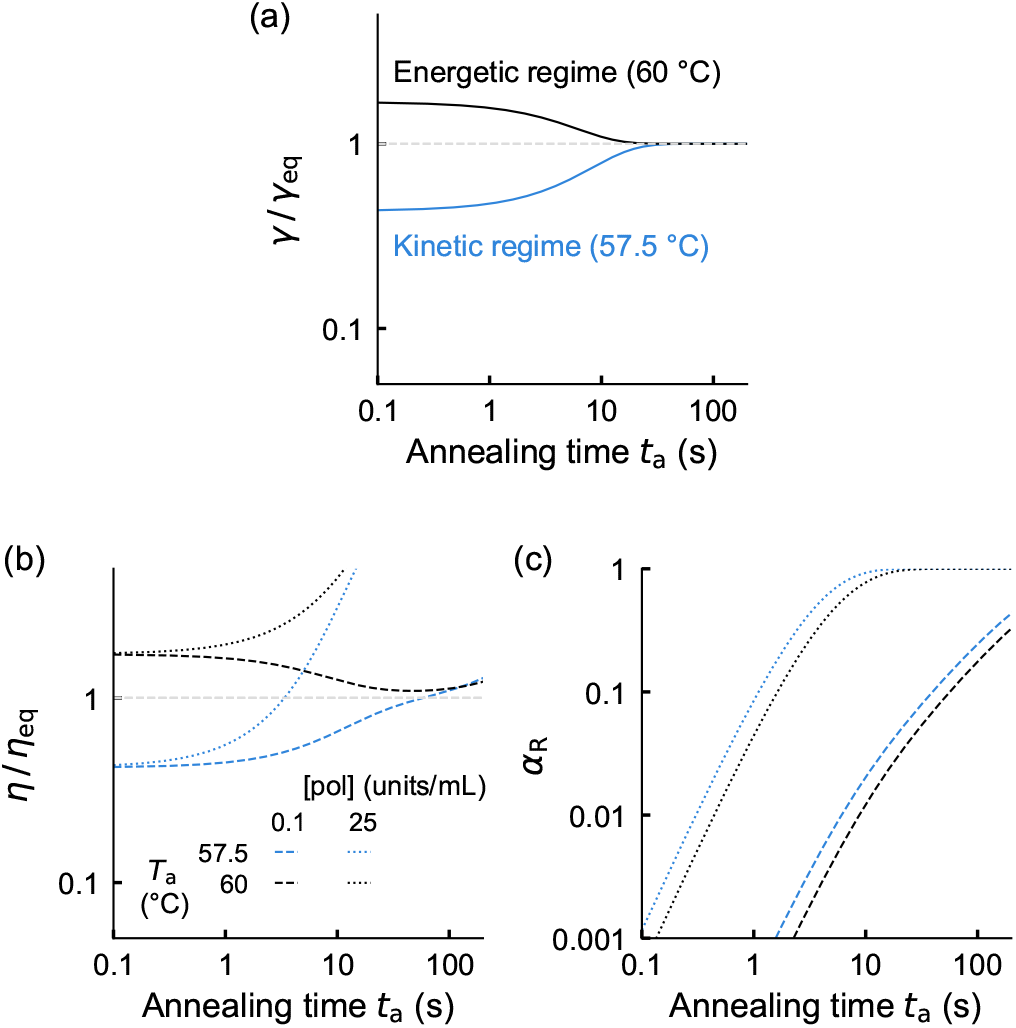
Dynamics of binding and error calculated by simulation. (a) Binding dynamics in the absence of polymerase. *γ* is the fraction of the complex of the primer and wrong template. *γ*_eq_ is its equilibrium value. See the main text. *γ*_eq_ = 0.038 for 60 °C and 0.057 for 57.5 °C. (b) Error rate dynamics in the presence of polymerase normalized by *η*_eq_ given by (2). The polymerase concentration is 0.1 units*/*mL (dashed) or 25 units*/*mL (dotted). *η*_eq_ = 0.0033 for 60 °C and 0.0045 for 57.5 °C. (c) The growth rates of the right product. For the simulation, the dissociation constants are obtained experimentally. 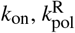 and 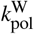 are obtained by the global fitting of the experimental results at each temperature. See SI Section S5 for details.

These binding dynamics imply that, in the presence of the polymerase, *η* may increase with *t*_a_ and be smaller than *η*_eq_ in the kinetic regime. That is, we may reduce errors and cycle time simultaneously and also achieve significant error suppression. This prediction was confirmed in the case of dilute polymerase (0.1 units*/*mL) and relatively short *t*_a_ (Fig. 5b). However, at *t*_a_ larger than ∼ 100 s, *η* increased with *t*_a_ in both regimes. This increase is pronounced with a typical polymerase concentration for PCR (25 units*/*mL). The increase in *η* at large *t*_a_ is caused by the saturation of the growth; *α*_W_ increases with *t*_a_ while *α*_R_ is already saturated (Fig. 5c). Nevertheless, we still obtained *η < η*_eq_ at small *t*_a_. At the limit of vanishing *t*_a_, *η* converges to

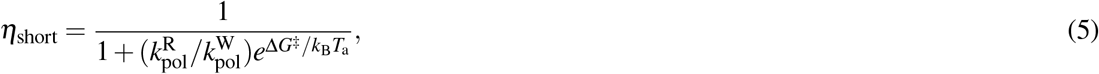

which has the same mathematical form as *η*_eq_ given by (2) but with ∆*G*^‡^ appearing instead of ∆*G* (SI Section S4).

It remains unclear whether the regime *η < η*_eq_ is experimentally accessible. The expected time scale *t*_a_ ∼ 1 s is significantly shorter than the typical time scale (∼ 10 s) necessary for cooling from ~ 95 °C at the denaturing step to ∼ 60 °C at the annealing step. Our model assumes an instantaneous temperature jump for simplicity and may no longer be applicable for such a short *t*_a_.

The short *t*_a_ results in a significant reduction of the product amount (Fig. 5c). However, the approach implementing the kinetic regime would be beneficial in the applications such as genome editing, where error suppression is critical.

### Multiple wrong sequences

In real-world applications of PCR, samples may contain multiple types of unwanted sequences. We study by numerical simulations of our model whether blockers could suppress replication errors in this case. For simplicity, we focus on a scenario in which the sample contains *N* types of wrong sequences, and we add *N* blocker sequences, each of which perfectly hybridizes to the corresponding wrong sequence. We fix the total concentration of the wrong sequences and the blocker sequences, so that the concentration of each wrong sequence and blocker sequence is proportional to 1*/N*.

We note that the concentration of B_W_: W is roughly proportional to [W][B_W_]. Since [W] and [B_W_] decrease with *N*, blocking may become less effective with *N*. The error rate *η* is defined similarly to (1), but where *α*_W_ is the total amount of wrong products (see SI section S4). We find that, although the error increases with *N*, the blocker strands suppress *η* even in the large *N* limit (Fig. 6). Moreover, our model predicts that the addition of blockers should not significantly affect *α*_R_ unless they strongly hybridize to R.

**Figure 6.**
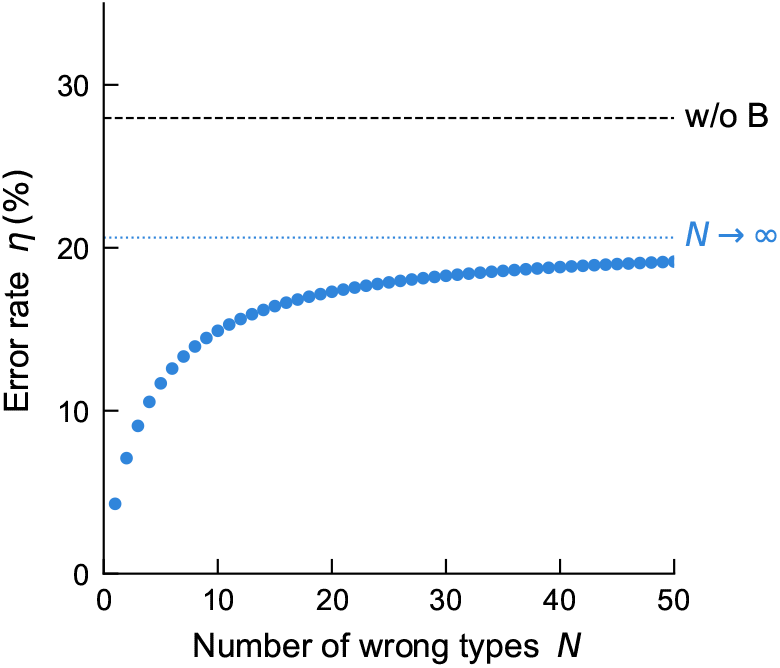
Error rate in the presence of multiple error sequences and blockers predicted by numerical simulation. The blue dotted line and black dashed line correspond to the result for a large number of wrong sequences (*N* = 10^4^) and without blocker strands, respectively.

## Discussion

Kinetic modeling of PCR reaction has contributed to quantitatively characterize the reaction performance^14–16^ and other aspects such as amplification heterogeneity^17^. However, modeling has been scarcely used to develop new guiding principles. The physics of information processing can provide such principles, thanks to its progress in characterizing general biochemical reactions^11,18–30^.

We demonstrated that adding blocker strands discriminates the right and wrong sequences by combining energetic and kinetic biasing. The kinetic biasing is effective in decreasing the error rate without affecting the efficiency. An alternative setup we studied is the use of blocker strands targeting the right template. In this case, we could increase the error rate up to a value larger than 80%. This inverted error control can not be achieved without kinetic biasing and may be helpful for sampling rare sequences.

Importantly, we found that error suppression is still effective at *T*_a_ much lower than *T*_m_ of the primer binding. This implies that we can suppress the hybridization errors in systems with limited temperature controllability, such as the reverse transcription, in which the optimal reaction temperature is not high, and the hybridization inside biological cells. This might also lead to a reduced cost of applications such as virus tests by using a low-cost cycler since we do not require accurate temperature control.

## Acknowledgements

ST was supported by JSPS KAKENHI Grant Numbers JP15H05460, JP18H05427, and JP19H01857. SP was supported by JSPS KAKENHI Grant Number JP18K03473 and by the Okawa Foundation (Grant Number 21-01).

## Methods

### Linear PCR experiment

DNA strands were synthesized by Eurofins Genomics, and Integrated DNA Technologies (see SI section S1). DNA/LNA chimeric strands were synthesized by Aji Bio-Pharma. The reaction mixture for polymerization contained Hot-start Taq DNA polymerase (New England Biolabs), Taq standard reaction buffer, R, W, P, and blocker strands. We performed initial heating for 30 s at 95 °C and, then, 10 or 40 cycles of 15 s at 95 °C, 30 s at 60 °C, and 5 s at 68 °C using a PCR cycler. Immediately after the cycles, the mixture was cooled down on the ice to stop the enzyme reaction and used for the quantification. The number of cycles were 40 when the polymerase concentration was 0.17, 0.25 or 0.5 units*/*mL and 10 otherwise.

### Quantification of P_R_ and P_W_

Additional quantitative PCR was performed on a real-time PCR cycler after the linear PCR experiment for quantifying *r* and *w* (see SI section S2). The reaction mixture contains Luna Universal qPCR Master Mix (New England Biolabs), 200 nM each of the primers, and the diluted sample. The dilution rate is 1/250 in the final concentration. The thermal cycle consists of initial heating for 60 s at 95 °C, 40 cycles of 15 s at 95 °C, 30 s at 66 °C, and 5 s at 72 °C.

### Melt curve analysis

We measured the melting curves for the hybridization of P, B_R_, and B_W_ to R and W from 95 °C and 20 °C and calculated their *K*_d_ (SI section S3). We mixed 100 nM each of DNA and double-strand-specific fluorescent molecule EvaGreen (Biotium). The fluorescent profile was analyzed based on the exponential background method^31^ to obtain the melting curve.

## Data availability

The data that support the findings of this study are available from the corresponding author upon reasonable request.

## Author Contributions

HA, SP, and ST designed the research, developed the theory, and wrote the paper. HA and SO did experiments.

## Competing Interests

The authors declare no conflict of interest.

## Notes

### Competing Interest Statement

The authors have declared no competing interest.

### Summary of Updates

Substantial changes were made, including the change in the main claim and the addition of new "Kinetic regime" section. Supplemental file was updated.

