## Supplementary texts, tables, and figures for "Error-suppression mechanism of PCR by blocker strands"

### Supplementary Methods and Results

#### S1 Linear PCR

**Sequences.** Sequences of the DNA strands used in the linear PCR step (Fig. S1) are presented in Table. S1. The sequences of the primer-binding regions of R and W differ by a single base, resulting in a single-base mismatch between P and W. The sequences outside the primer-binding region of R and W are different. The primer P includes a tag sequence at its 5’ end for the subsequent quantification step. The polymerase elongates the primers of the hybridized complexes P:R and P:W to produce $\overline{\mathrm{R}}$ and $\overline{\mathrm{W}}$, respectively. The 3’ ends of R and W include two-base overhang to prevent elongation from these ends.

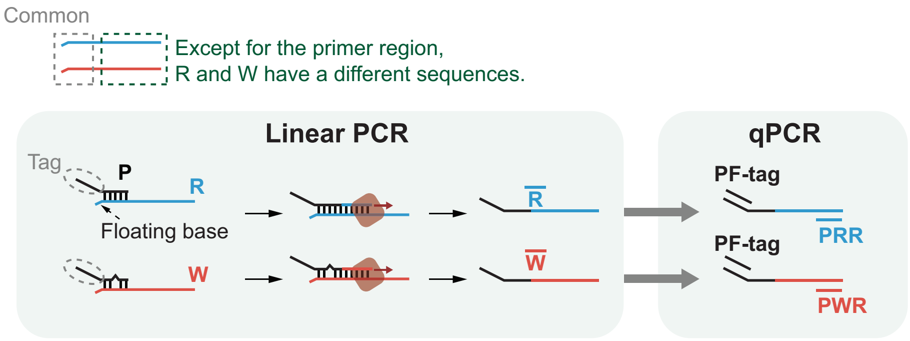


**Figure S1.** Scheme of the experiments. R and W have similar sequences in their primer region and are otherwise different. P includes a tag sequence for the subsequent quantitative PCR (qPCR). Two bases at the 3’ ends of R and W are floating to prevent elongation of the primer tag sequence.

The blockers $\mathrm{B_W}$ and $\mathrm{B_R}$ are LNA-DNA chimeric strands. They hybridize to a part of the primer-binding regions of R and W. As the blockers, we also used non-chimeric DNA strands which have the same sequence as $\mathrm{B_W}$ and $\mathrm{B_R}$ but do not contain LNA bases (Fig. S7).

**Time course.** We verified that the products grow linearly with cycles, indicating that the primer strands do not deplete under our experimental conditions (Fig. S2). In the experiments, we stopped reactions after the indicated number of cycles by immediately transferring the samples to on-ice environment.

**Table S1.** The DNA sequences used in the linear PCR step. LNA bases are indicated with lowercase.

| Name | Sequence |
|---|---|
| R | 5’-CTCTCGTTCTCAACCTACCATGGCAAGGGTTGTTCTTTGTTGGTGTCCACTGAC GACGC A TATCTCAAGGTA-3’ |
| W | 5’-TACATATGACGCACAGATGCAGCCAACTCCACTCACTTCTCTCTTCAGATTGAC GACGC T TATCTCAAGGTA-3’ |
| $\overline{R}$ | 5’-GTCACCCAATCGTCCTAGATCCTTGAGATATGCGTCGTCAGTGGACACCAACAA AGAACAACCCTTGCCATGGTAGGTTGAGAACGAGAG-3’ |
| $\overline{W}$ | 5’-GTCACCCAATCGTCCTAGATCCTTGAGATATGCGTCGTCAATCTGAAGAGAGAA GTGAGTGGAGTTGGCTGCATCTGTGCGTCATATGTA-3’ |
| P | 5’-GTCACCCAATCGTCCTAGATCCTTGAGATATGCGTCGTCA-3’ |
| $B_W$ | 5’-GAGATa a gcGTCGTTT-3’ |
| $B_R$ | 5’-GAGATa t gcGTCGTTT-3’ |
| $B_W$ (DNA) | 5’-GAGATA A GCGTCGTTT-3’ |
| $B_R$ (DNA) | 5’-GAGATA T GCGTCGTTT-3’ |

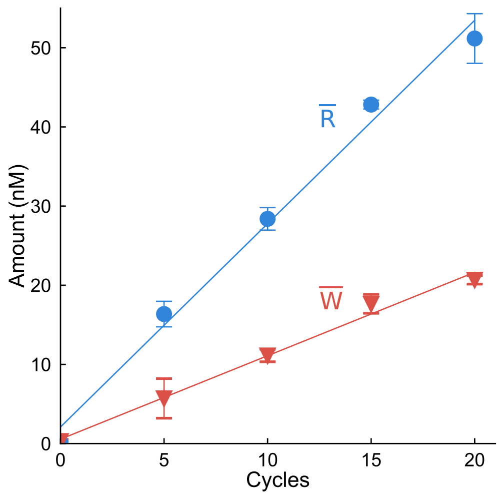


**Figure S2.** Time course of the polymerase reaction. The products increase linearly during the cycles. The error bars represent standard deviations.

### S2 Quantitative PCR

**Sequences.** We use two primer sets {PF-tag, PRR} and {PF-tag, PWR} (Table. S2) for detecting $\overline{\mathrm{R}}$ and $\overline{\mathrm{W}}$, respectively. The PRR and PWR hybridize to the 21-base region at the 3' end of $\overline{\mathrm{R}}$ and $\overline{\mathrm{W}}$, respectively. The reaction mixture contains diluted sample of the linear PCR (diluted to 1/250 in the final concentration), 200-nM each primers and Luna Universal qPCR Master Mix (New England Biolabs).

**Table S2.** The DNA sequences for the quantitative PCR.

| Name | Sequence |
|---|---|
| PF-tag | 5'-GTCACCCAATCGTCCTAGATC-3' |
| PRR | 5'-CTCTCGTTCTCAACCTACCAT-3' |
| PWR | 5'-TACATATGACGCACAGATGCA-3' |

**Standard curves.** We measured the standard curves used for the quantification of $\overline{\mathrm{R}}$ and $\overline{\mathrm{W}}$ (Fig. S3). The residues of P, R, and W used in the linear PCR step remain in the subsequent quantitative PCR step. Although they are diluted to 1/250, they increase the background level and affect the quantification when the template concentration is low. Therefore, for making the standard curves, we added 400 pM P and 6.25 pM R and 6.25 pM W to imitate the effect of the residues. In addition to these strands, the reaction mixture contains Luna Universal qPCR Master Mix (New England Biolabs), templates $\overline{\mathrm{R}}$ or $\overline{\mathrm{W}}$, and 200-nM each primer sets, {PF-tag, PRR} and {PF-tag, PWR}. The template concentrations were 160, 20, 2.5, 0.31, 0.039, and 0.049 pM.

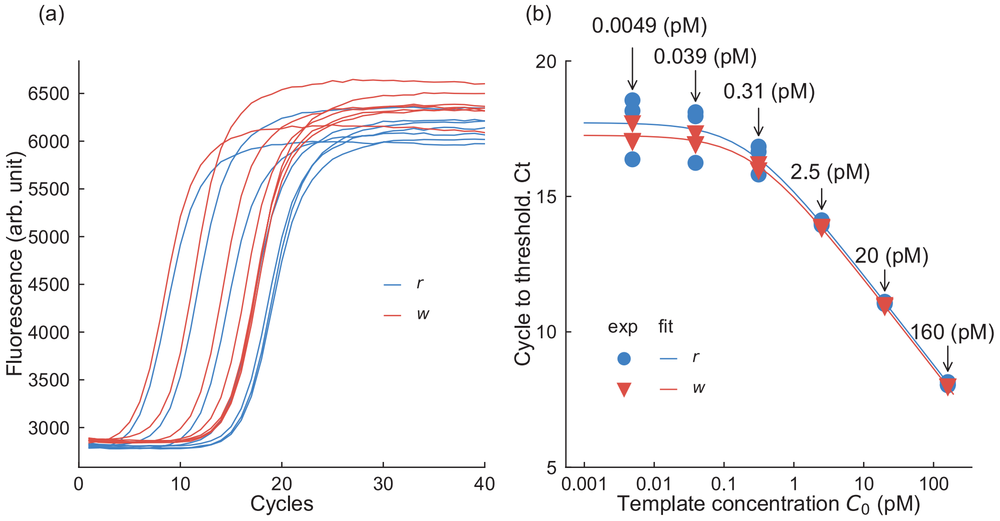


**Figure S3.** Standard curve measurement. (a) The qPCR curves with different initial template concentrations. (b) Standard curves. $\mathrm{C_t}$ values saturate at low template concentrations due to mainly the contamination of the residues of the linear PCR step. We directly fitted these points with Eq. (S1) with $\delta$ and $\phi$ as the fitting parameters (solid curves).

The $\mathrm{C_t}$ values, which are defined by the cycle number where the fluorescence reached a threshold value, are related to the template concentration $C_0$ by

$$\mathrm{C_t} = -\log_2[C_0 + \delta] + \phi. \tag{S1}$$

We calculated $\mathrm{C_t}$ based on the maxRatio method[1]. We fitted the experimental data by this relation (Fig. S3b) and obtained the background level $\delta$ (15.4 for R and 15.3 for W) and intercept $\phi$ (0.204 for R and 0.256 for W). Using these parameters, $[\overline{\mathrm{R}}]$ and $[\overline{\mathrm{W}}]$ are calculated from $\mathrm{C_t}$ values by

$$[\overline{\mathrm{R}}] = 2^{-(\mathrm{C_t}-15.4)} - 0.204,$$
$$[\overline{\mathrm{W}}] = 2^{-(\mathrm{C_t}-15.3)} - 0.256. \tag{S2}$$

### S3 Melting curve analysis for measuring $T_{\rm m}$, $\Delta H^{\rm o}$, $\Delta S^{\rm o}$, and $K_{\rm d}$

We evaluated the melting temperatures $T_{\rm m}$ and dissociation constants $K_{\rm d}$ by a melting curve analysis.

**Sequences.** We used shortened strands pR, pW, and pP instead of R, W, and P, respectively, to eliminate the effect of the secondary structures (Table. S3). The sequences of pR and pW correspond to the primer binding regions of R and W, respectively. The sequence of pP is the same as P, but without the tag sequence.

**Table S3.** The DNA sequences that we used for the melting curve analysis.

| Name | Sequence |
|---|---|
| pR | 5'-TGACGACGCATATCTCAAGGTA-3' |
| pW | 5'-TGACGACGCTTATCTCAAGGT-3' |
| pP | 5'-ATCCTTGAGATATGCGTCGTCA-3' |

**Analysis.** We normalized the fluorescence intensity by the exponential background method[2] (Fig. S4), which provides the normalized fraction $a$ of the DNA strands that are forming double strands. The dependence of $\Delta G^{\rm o}$ on the temperature is calculated from $a$ as

$$\Delta G^{\rm o} = RT \ln \left[ \frac{C_0(1-a)^2}{a} \right]. \tag{S3}$$

Here, $R$ is the gas constant, $T$ is the temperature, and $C_0 = 100\,\text{nM}$ is the concentration of each DNA. On the other hand, we have that

$$\Delta G^{\rm o} = \Delta H^{\rm o} - T\Delta S^{\rm o}. \tag{S4}$$

Thus, we could obtain $\Delta H^{\rm o}$ and $\Delta S^{\rm o}$ by fitting the $\Delta G^{\rm o}$ between the melting region with Eq. S4. From the The obtained $T_{\rm m}$ and $\Delta H^{\rm o}$, $\Delta S^{\rm o}$, and $K_{\rm d}$ are shown in Table. S4. These thermodynamic parameters allow us to obtain the standard free energy change $\Delta G^{\rm o}$ at arbitrary temperature, assuming that $\Delta H^{\rm o}$ and $\Delta S^{\rm o}$ are temperature independent[3]. $T_{\rm m}$ values reflecting DNA concentrations in our experiments are obtained by,

$$T_{\rm m} = \frac{\Delta H^{\rm o}}{\Delta S^{\rm o} + R\ln([\text{ss1}] - [\text{ss2}]/2)}. \tag{S5}$$

Here, $[\text{ss1}]$ is concentration and larger than that of the complementary sequence $[\text{ss2}]$.

**Table S4.** Melting temperature $T_{\rm m}$ is calculated as the temperature that is calculated by Eq. S5. $K_{\rm d}$ at 60 °C and 57.5 °C are calculated from Eq. S4.

| Pair | $T_{\rm m}$ (°C) | $\Delta H^{\rm o}$ (kcal/mol) | $\Delta S^{\rm o}$ (cal/mol) | $K_{\rm d}$ at 60 °C (nM) | $K_{\rm d}$ at 57.5 °C (nM) |
|---|---|---|---|---|---|
| P : R | 62.8 | -163.4 | 454.3 | 12.43 | 1.922 |
| P : W | 59.3 | -160.6 | -451.0 | 162.2 | 25.91 |
| $B_{\rm R}$ : R | 65.5 | -119.6 | -327.1 | 106.7 | 27.22 |
| $B_{\rm R}$ : W | 57.5 | -98.36 | -271.4 | 6149 | 1999 |
| $B_{\rm W}$ : R | 56.3 | -100.5 | -279.0 | 11120 | 3527 |
| $B_{\rm W}$ : W | 65.3 | -117.5 | -321.1 | 124.3 | 32.49 |

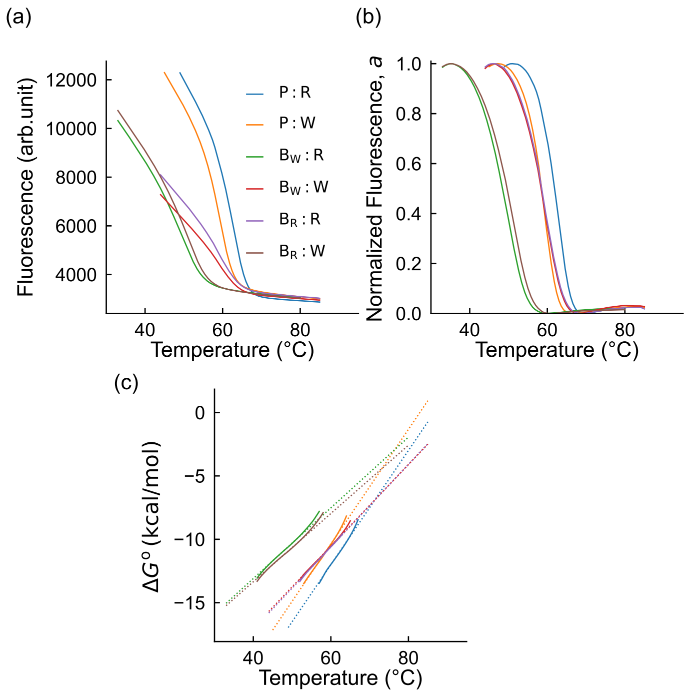


**Figure S4.** Melting curves. (a) Fluorescence curves. (b) Normalized fluorescence curves obtained by the exponential background method[2]. (c) The solid lines are the $\Delta G^{\mathrm{o}}$ caluculated by the Eq. S3. The dotted lines are the results of the fitting with Eq. S4.

### S4 Theoretical analysis

We consider, in general, a system containing $N$ types of wrong sequences $W_i$, with $i = 1 \dots N$. We model the hybridization dynamics during the annealing step by the rate equations

$$
\begin{aligned}
\frac{d[P:R]}{dt} &= k_{on}[R]_f[P]_f - k_{off}^{P:R}[P:R] - k_{pol}^{R}[pol][P:R],\\
\frac{d[\overline{R}:R]}{dt} &= k_{pol}^{R}[pol][P:R],\\
\frac{d[P:W_i]}{dt} &= k_{on}[W_i]_f[P]_f - k_{off}^{P:W_i}[P:W_i] - k_{pol}^{W_i}[pol][P:W_i],\\
\frac{d[\overline{W}_i:W_i]}{dt} &= k_{pol}^{W_i}[pol][P:W_i],\\
\frac{d[B_j:R]}{dt} &= k_{on}[R]_f[B_j]_f - k_{off}^{B_j:R}[B_j:R],\\
\frac{d[B_j:W_i]}{dt} &= k_{on}[W_i]_f[B_j]_f - k_{off}^{B_j:W_i}[B_j:W_i].
\end{aligned}
\tag{S6}
$$

Here, $[\cdot]$ denotes concentration of single DNA strands and $[\cdot]_f$ denotes the concentration of single DNA strands that are not hybridizing to other strands. The conservation laws read

$$
\begin{aligned}
[P] &= [P]_f + [P:R] + \sum_i^N [P:W_i] + [\overline{R}:R] + \sum_i^N [\overline{W}_i:W_i],\\
[R] &= [R]_f + [P:R] + \sum_j^N [B_j:R] + [\overline{R}:R],\\
[W_i] &= [W_i]_f + [P:W_i] + \sum_j^N [B_j:W_i] + [\overline{W}_i:W_i],\\
[B_j] &= [B_j]_f + \sum_i^N [B_j:W_i] + [B_j:R].
\end{aligned}
\tag{S7}
$$

When $[B] \gg [P]$, we can assume that the last two equations of (S6) are stationary, i.e., $d[B_j:R]/dt = d[B_j:W_i]/dt = 0$.

$$
[R:B_i] = \frac{k_{on}[R]_f[B_i]_f}{k_{off}^{B_i:R}} = \frac{[R]_f[B_i]_f}{K_d^{B_i:R}}, \qquad [B_j:W_i] = \frac{k_{on}[W_i]_f[B_j]_f}{k_{off}^{B_j:W_i}} = \frac{[W_i]_f[B_j]_f}{K_d^{B_j:W_i}}. \tag{S8}
$$

This assumption holds well in our experiments with $[B] = 2000\,\mathrm{nM}$ and $[P] = 100\,\mathrm{nM}$. Since $[B],[P] \gg [R] = [W] = 2.5\,\mathrm{nM}$, we also make the approximation

$$
[P]_f \simeq [P], \qquad [B_i]_f \simeq [B_i]. \tag{S9}
$$

By substituting Eq. S8 and Eq. S9 to Eq. S6, we obtain

$$
\begin{aligned}
\frac{d[P:R]}{dt} &= k_{on,eff}^{R}[P]\left([R] - [P:R] - [\overline{R}:R]\right) - k_{off,eff}^{P:R}[P:R]\\
\frac{d[P:W_i]}{dt} &= k_{on,eff}^{W_i}[P]\left([W_i] - [P:W_i] - [\overline{W}_i:W_i]\right) - k_{off,eff}^{P:W_i}[P:W_i],
\end{aligned}
\tag{S10}
$$

where we defined the effective rate constants

$$
k_{on,eff}^{R} = \frac{k_{on}}{1+\sum_j \frac{[B_j]}{K_d^{B_j:R}}}, \quad k_{on,eff}^{W_i} = \frac{k_{on}}{1+\sum_j \frac{[B_j]}{K_d^{B_j:W_i}}}, \quad k_{off,eff}^{P:R} = k_{off}^{P:R} + k_{pol}^{R}[pol], \quad k_{off,eff}^{P:W_i} = k_{off}^{P:W_i} + k_{pol}^{W_i}[pol]. \tag{S11}
$$

By solving (S10), we obtain

$$
\begin{aligned}
[P:R] &= \frac{k_{on,eff}^{R}[P][R]}{\lambda_{+,R} - \lambda_{-,R}}\left(e^{\lambda_{+,R}t} - e^{\lambda_{-,R}t}\right),\\
[P:W_i] &= \frac{k_{on,eff}^{W_i}[P][W_i]}{\lambda_{+,W_i} - \lambda_{-,W_i}}\left(e^{\lambda_{+,W_i}t} - e^{\lambda_{-,W_i}t}\right).
\end{aligned}
\tag{S12}
$$

Furthermore, by substituting these equations to second and fourth equations in (S6), we obtain

$$
\begin{aligned}
[\overline{\mathrm{R}}:\mathrm{R}] &= [\mathrm{R}]\left(1+\frac{\lambda_{-,\mathrm{R}}}{\lambda_{+,\mathrm{R}}-\lambda_{-,\mathrm{R}}}e^{\lambda_{+,\mathrm{R}}t}-\frac{\lambda_{+,\mathrm{R}}}{\lambda_{+,\mathrm{R}}-\lambda_{-,\mathrm{R}}}e^{\lambda_{-,\mathrm{R}}t}\right),\\
[\overline{\mathrm{W}}_i:\mathrm{W}_i] &= [\mathrm{W}_i]\left(1+\frac{\lambda_{-,\mathrm{W}_i}}{\lambda_{+,\mathrm{W}_i}-\lambda_{-,\mathrm{W}_i}}e^{\lambda_{+,\mathrm{W}_i}t}-\frac{\lambda_{+,\mathrm{W}_i}}{\lambda_{+,\mathrm{W}_i}-\lambda_{-,\mathrm{W}_i}}e^{\lambda_{-,\mathrm{W}_i}t}\right),
\end{aligned}
\tag{S13}
$$

where

$$
\begin{aligned}
\lambda_{\pm,\mathrm{R}} &= -\frac{1}{2}\left(k^{\mathrm{R}}_{\mathrm{on,eff}}[\mathrm{P}]+k^{\mathrm{P:R}}_{\mathrm{off,eff}}\right)\pm\frac{1}{2}\sqrt{\left(k^{\mathrm{R}}_{\mathrm{on,eff}}[\mathrm{P}]+k^{\mathrm{P:R}}_{\mathrm{off,eff}}\right)^2-4k^{\mathrm{R}}_{\mathrm{on,eff}}[\mathrm{P}]k_{\mathrm{pol}}[\mathrm{pol}]},\\
\lambda_{\pm,\mathrm{W}_i} &= -\frac{1}{2}\left(k^{\mathrm{W}_i}_{\mathrm{on,eff}}[\mathrm{P}]+k^{\mathrm{P:W}_i}_{\mathrm{off,eff}}\right)\pm\frac{1}{2}\sqrt{\left(k^{\mathrm{W}_i}_{\mathrm{on,eff}}[\mathrm{P}]+k^{\mathrm{P:W}_i}_{\mathrm{off,eff}}\right)^2-4k^{\mathrm{W}_i}_{\mathrm{on,eff}}[\mathrm{P}]k_{\mathrm{pol}}[\mathrm{pol}]}.
\end{aligned}
\tag{S14}
$$

**The barrier height difference $\Delta G^{\ddagger}$.** Eq. (S11) implies that the addition of the blocker alters the activation energy to form the hybridization. In fact, assuming the Arrhenius description, the barrier heights are expressed as follows.

$$
G^{\ddagger}_{\mathrm{R}} = RT_{\mathrm{a}}\ln\frac{A}{k_{\mathrm{on}}}+k_{\mathrm{B}}T_{\mathrm{a}}\ln\left[1+\sum_j\frac{[\mathrm{B}_j]}{K^{\mathrm{B}_j:\mathrm{R}}_{\mathrm{d}}}\right],\quad G^{\ddagger}_{\mathrm{W}_i} = k_{\mathrm{B}}T_{\mathrm{a}}\ln\frac{A}{k_{\mathrm{on}}}+k_{\mathrm{B}}T_{\mathrm{a}}\ln\left[1+\sum_j\frac{[\mathrm{B}_j]}{K^{\mathrm{B}_j:\mathrm{W}_i}_{\mathrm{d}}}\right].
\tag{S15}
$$

Here, $A$ is the collision factor. Thus,

$$
\Delta G^{\ddagger} = G^{\ddagger}_{\mathrm{W}_i}-G^{\ddagger}_{\mathrm{R}} \simeq k_{\mathrm{B}}T_{\mathrm{a}}\ln\frac{1+\sum_j[\mathrm{B}_j]/K^{\mathrm{B}_j:\mathrm{W}_i}_{\mathrm{d}}}{1+\sum_j[\mathrm{B}_j]/K^{\mathrm{B}_j:\mathrm{R}}_{\mathrm{d}}}.
\tag{S16}
$$

**Free energy difference of the hybridized states.** Next, we evaluate the free energy difference between P : R and P : $\mathrm{W_i}$. By solving (S10) with $[\mathrm{pol}] = 0$ to focus on the hybridization dynamics, we otain

$$
[\mathrm{P}:\mathrm{R}] = [\mathrm{P}:\mathrm{R}]_{\mathrm{eq}}\left(1-e^{-t/\tau_{\mathrm{P:R}}}\right),\qquad [\mathrm{P}:\mathrm{W}_i] = [\mathrm{P}:\mathrm{W}_i]_{\mathrm{eq}}\left(1-e^{-t/\tau_{\mathrm{P:W}_i}}\right),
\tag{S17}
$$

where

$$
\begin{aligned}
\tau_{\mathrm{P:R}} &= \left(\frac{k_{\mathrm{on}}[\mathrm{P}]}{1+\sum_j^N\frac{[\mathrm{B}_j]}{K^{\mathrm{B}_j:\mathrm{R}}_{\mathrm{d}}}}+k^{\mathrm{P:R}}_{\mathrm{off}}\right)^{-1},\quad [\mathrm{P}:\mathrm{R}]_{\mathrm{eq}} = \frac{[\mathrm{R}][\mathrm{P}]}{1+\sum_j^N\frac{[\mathrm{B}_j]}{K^{\mathrm{B}_j:\mathrm{R}}_{\mathrm{d}}}}\left(\frac{[\mathrm{P}]}{1+\sum_j^N\frac{[\mathrm{B}_j]}{K^{\mathrm{B}_j:\mathrm{R}}_{\mathrm{d}}}}+K^{\mathrm{P:R}}_{\mathrm{d}}\right)^{-1},\\
\tau_{\mathrm{P:W}_i} &= \left(\frac{k_{\mathrm{on}}[\mathrm{P}]}{1+\sum_j^N\frac{[\mathrm{B}_j]}{K^{\mathrm{B}_j:\mathrm{W}_i}_{\mathrm{d}}}}+k^{\mathrm{P:W}_i}_{\mathrm{off}}\right)^{-1},\quad [\mathrm{P}:\mathrm{W}_i]_{\mathrm{eq}} = \frac{[\mathrm{W}_i][\mathrm{P}]}{1+\sum_j^N\frac{[\mathrm{B}_j]}{K^{\mathrm{B}_j:\mathrm{W}_i}_{\mathrm{d}}}}\left(\frac{[\mathrm{P}]}{1+\sum_j^N\frac{[\mathrm{B}_j]}{K^{\mathrm{B}_j:\mathrm{W}_i}_{\mathrm{d}}}}+K^{\mathrm{P:W}_i}_{\mathrm{d}}\right)^{-1}.
\end{aligned}
\tag{S18}
$$

Since the fraction of the equilibrium concentration is related to the free energy difference $\Delta G$, we obtain

$$
\mathrm{e}^{-\Delta G/RT_{\mathrm{a}}} = \frac{[\mathrm{P}:\mathrm{W}_i]_{\mathrm{eq}}}{[\mathrm{P}:\mathrm{R}]_{\mathrm{eq}}} \approx \frac{[\mathrm{W}_i]\left[1+\frac{K^{\mathrm{P:R}}_{\mathrm{d}}}{[\mathrm{P}]}\left(1+\sum_j\frac{[\mathrm{B}_j]}{K^{\mathrm{B}_j:\mathrm{R}}_{\mathrm{d}}}\right)\right]}{[\mathrm{R}]\left[1+\frac{K^{\mathrm{P:W}_i}_{\mathrm{d}}}{[\mathrm{P}]}\left(1+\sum_j\frac{[\mathrm{B}_j]}{K^{\mathrm{B}_j:\mathrm{W}_i}_{\mathrm{d}}}\right)\right]}.
\tag{S19}
$$

**Comparison between $\Delta G^{\ddagger}$ and $\Delta G$.** In our experiments, we have only one type of wrong sequence W and a single blocker species. In the presence of $\mathrm{B_W}$, we obtain

$$
\begin{aligned}
\Delta G^{\ddagger} &\approx k_{\mathrm{B}}T_{\mathrm{a}}\ln\left[\frac{1+[\mathrm{B_W}]/K^{\mathrm{B_W:W}}_{\mathrm{d}}}{1+[\mathrm{B_W}]/K^{\mathrm{B_W:R}}_{\mathrm{d}}}\right]\approx k_{\mathrm{B}}T_{\mathrm{a}}\ln\left(1+\frac{[\mathrm{B_W}]}{K^{\mathrm{B_W:W}}_{\mathrm{d}}}\right),\\
\Delta G &\approx k_{\mathrm{B}}T_{\mathrm{a}}\ln\left[\frac{1+\frac{K^{\mathrm{P:W}}_{\mathrm{d}}}{[\mathrm{P}]}\left(1+\frac{[\mathrm{B_W}]}{K^{\mathrm{B_W:W}}_{\mathrm{d}}}\right)}{1+\frac{K^{\mathrm{P:R}}_{\mathrm{d}}}{[\mathrm{P}]}\left(1+\frac{[\mathrm{B_W}]}{K^{\mathrm{B_W:R}}_{\mathrm{d}}}\right)}\right]\approx k_{\mathrm{B}}T_{\mathrm{a}}\ln\left[1+\frac{K^{\mathrm{P:W}}_{\mathrm{d}}}{[\mathrm{P}]}\left(1+\frac{[\mathrm{B_W}]}{K^{\mathrm{B_W:W}}_{\mathrm{d}}}\right)\right],\\
\Delta G_0 &\approx k_{\mathrm{B}}T_{\mathrm{a}}\ln\left[\frac{1+\frac{K^{\mathrm{P:W}}_{\mathrm{d}}}{[\mathrm{P}]}}{1+\frac{K^{\mathrm{P:R}}_{\mathrm{d}}}{[\mathrm{P}]}}\right]\approx k_{\mathrm{B}}T_{\mathrm{a}}\ln\left[1+\frac{K^{\mathrm{P:W}}_{\mathrm{d}}}{[\mathrm{P}]}\right].
\end{aligned}
\tag{S20}
$$

It follows from these relations that

$$\Delta G - \Delta G_0 \approx k_B T_a \ln \left( 1 + \frac{1}{1+[P]/K_d^{P:W}} \frac{[B_W]}{K_d^{B_W:W}} \right). \tag{S21}$$

**S4.1. Limit of fast polymerase reaction.** Here, we assume that the elongation reaction is so fast and take place before the primer dissociates from the template strand. Under this assumption, it is reasonable to ignore the dissociation terms of the first and third equations in Eq. S6. The effective off-rate constant would be $k_{\mathrm{off,eff}}^{\mathrm{P:R}} \approx k_{\mathrm{pol}}^{\mathrm{R}}[\mathrm{pol}]$, $k_{\mathrm{off,eff}}^{\mathrm{P:W}} \approx k_{\mathrm{pol}}^{\mathrm{W}}[\mathrm{pol}]$. Taking the limit of the polymerase concentration, Eq. S13 becomes,

$$\begin{aligned} [\overline{\mathrm{R}}:\mathrm{R}] &\approx 1 - e^{-k_{\mathrm{on,eff}}^{\mathrm{R}}[\mathrm{P}]t}, \\ [\overline{\mathrm{W}}:\mathrm{W}] &\approx 1 - e^{-k_{\mathrm{on,eff}}^{\mathrm{W}}[\mathrm{P}]t}. \end{aligned} \tag{S22}$$

The error rate is

$$\eta = \frac{[\overline{\mathrm{W}}:\mathrm{W}]}{[\overline{\mathrm{R}}:\mathrm{R}]+[\overline{\mathrm{W}}:\mathrm{W}]} \approx \frac{1}{1+\frac{1-\exp(-k_{\mathrm{on,eff}}^{\mathrm{R}}[\mathrm{P}]t)}{1-\exp(-k_{\mathrm{on,eff}}^{\mathrm{W}}[\mathrm{P}]t)}} \tag{S23}$$

When the blocker is $\mathrm{B_W}$, $k_{\mathrm{on,eff}}^{\mathrm{R}}[\mathrm{P}]t$ become large, and $k_{\mathrm{on,eff}}^{\mathrm{W}}[\mathrm{P}]t$ become small. The error rate becomes,

$$\eta_{\mathrm{fast}} \approx \frac{1}{1+(k_{\mathrm{on}}[\mathrm{P}]t)^{-1}\exp(\Delta G^{\ddagger}/T_a)} \tag{S24}$$

**Limit of slow polymerase reaction.** Here, we assume that the elongation reaction does not deplete the hybridized products [P : R] and [P : W]. Under this assumption, it becomes reasonable to ignore the elongation terms of the first and third equations in Eq. S6. By solving them again, we obtain,

$$\begin{aligned} [\overline{\mathrm{R}}:\mathrm{R}] &= k_{\mathrm{pol}}^{\mathrm{R}}[\mathrm{pol}][\mathrm{P}:\mathrm{R}]_{\mathrm{eq}} \left( t + \tau_{\mathrm{P:R}} e^{-t/\tau_{\mathrm{P:R}}} - \tau_{\mathrm{P:R}} \right), \\ [\overline{\mathrm{W}}:\mathrm{W}] &= k_{\mathrm{pol}}^{\mathrm{W}}[\mathrm{pol}][\mathrm{P}:\mathrm{W}]_{\mathrm{eq}} \left( t + \tau_{\mathrm{P:W}} e^{-t/\tau_{\mathrm{P:W}}} - \tau_{\mathrm{P:W}} \right). \end{aligned} \tag{S25}$$

The error rate is,

$$\eta = \frac{[\overline{\mathrm{W}}:\mathrm{W}]}{[\overline{\mathrm{R}}:\mathrm{R}]+[\overline{\mathrm{W}}:\mathrm{W}]} = \frac{1}{1+\frac{k_{\mathrm{pol}}^{\mathrm{R}}}{k_{\mathrm{pol}}^{\mathrm{W}}} e^{\Delta G/T_a} \frac{t+\tau_{\mathrm{P:R}}e^{-t/\tau_{\mathrm{P:R}}}-\tau_{\mathrm{P:R}}}{t+\tau_{\mathrm{P:W}}e^{-t/\tau_{\mathrm{P:W}}}-\tau_{\mathrm{P:W}}}}. \tag{S26}$$

If the polymerase concentration is sufficiently low, we are able to assume that the template DNA does not deplete even when $t \to \infty$. Then the error rate is expressed as,

$$\lim_{t\to\infty} \eta = \eta_{\mathrm{eq}} = \frac{1}{1+\frac{k_{\mathrm{pol}}^{\mathrm{R}}}{k_{\mathrm{pol}}^{\mathrm{W}}} e^{\Delta G/k_B T_a}}. \tag{S27}$$

**Short time limit.** At the vanishing limit of the annealing time, Taylor's expansion of Eq. S13 gives,

$$\begin{aligned} [\overline{\mathrm{R}}:\mathrm{R}] &= [\mathrm{R}]\lambda_+^{\mathrm{R}}\lambda_-^{\mathrm{R}}t^2 = k_{\mathrm{on,eff}}^{\mathrm{R}}k_{\mathrm{pol}}^{\mathrm{R}}[\mathrm{P}][\mathrm{R}][\mathrm{pol}], \\ [\overline{\mathrm{W}}:\mathrm{W}] &= [\mathrm{W}]\lambda_+^{\mathrm{W}}\lambda_-^{\mathrm{W}}t^2 = k_{\mathrm{on,eff}}^{\mathrm{W}}k_{\mathrm{pol}}^{\mathrm{W}}[\mathrm{P}][\mathrm{W}][\mathrm{pol}]. \end{aligned} \tag{S28}$$

The error rate is,

$$\eta_{\mathrm{short}} = \frac{[\overline{\mathrm{W}}:\mathrm{W}]}{[\overline{\mathrm{R}}:\mathrm{R}]+[\overline{\mathrm{W}}:\mathrm{W}]} = \frac{1}{1+(k_{\mathrm{pol}}^{R}/k_{\mathrm{pol}}^{\mathrm{W}})e^{\Delta G^{\ddagger}/k_B T_a}}. \tag{S29}$$

**Multiple errors.** For deriving the results in Fig. 5, we numerically solved the rate equations (S6) in the presence of multiple types of wrong sequences and corresponding blocker sequences. We fixed the total concentration of the wrong-template sequences $[\mathrm{W}] \equiv \sum_i^N [\mathrm{W}_i]$ and the blocker sequences $[\mathrm{B}] \equiv \sum_i^N [\mathrm{B}_i]$ and changed $N$ to see how the the number of species $N$ affect the error rate $\eta$. The wrong-template and blocker concentrations of each species are assumed to be the same; $[\mathrm{W}_i] \equiv [\mathrm{W}]$ and $[\mathrm{B}_i] \equiv [\mathrm{B}]$. Accordingly, $k_{\mathrm{off}}$ for $\mathrm{B_j:W_i}$ take two values depending on whether the hybridization is specific or has a mismatch; $k_{\mathrm{off}}^{\mathrm{B:W,spc}}$ if $i=j$ and $k_{\mathrm{off}}^{\mathrm{B:W,mis}}$ if $i \neq j$. $k_{\mathrm{off}}^{\mathrm{B}_j\mathrm{:R}}$ are independent of $j$.

Following these assumptions, the equations we used for the simulations are

$$
\begin{aligned}
\frac{\mathrm{d}[\mathrm{P:R}]}{\mathrm{d}t} &= k_{\mathrm{on}}[\mathrm{P}]_{\mathrm{f}}[\mathrm{R}]_{\mathrm{f}} - k_{\mathrm{off}}^{\mathrm{P:R}}[\mathrm{P:R}] - k_{\mathrm{pol}}^{\mathrm{R}}[\mathrm{pol}][\mathrm{P:R}],\\
\frac{\mathrm{d}[\overline{\mathrm{R}}:\mathrm{R}]}{\mathrm{d}t} &= k_{\mathrm{pol}}^{\mathrm{R}}[\mathrm{pol}][\mathrm{P:R}],\\
\frac{\mathrm{d}[\mathrm{P:W}]}{\mathrm{d}t} &= Nk_{\mathrm{on}}[\mathrm{P}]_{\mathrm{f}}[\mathrm{W}]_{\mathrm{f}} - k_{\mathrm{off}}^{\mathrm{P:W}_i}[\mathrm{P:W}] - k_{\mathrm{pol}}^{\mathrm{W}}[\mathrm{pol}][\mathrm{P:W}],\\
\frac{\mathrm{d}[\overline{\mathrm{W}}:\mathrm{W}]}{\mathrm{d}t} &= k_{\mathrm{pol}}^{\mathrm{W}}[\mathrm{pol}][\mathrm{P:W}],\\
\frac{\mathrm{d}[\mathrm{B:R}]}{\mathrm{d}t} &= Nk_{\mathrm{on}}[\mathrm{B}]_{\mathrm{f}}[\mathrm{R}]_{\mathrm{f}} - k_{\mathrm{off}}^{\mathrm{B:R}}[\mathrm{R:B}],\\
\frac{\mathrm{d}[\mathrm{B:W}]_{\mathrm{mis}}}{\mathrm{d}t} &= N(N-1)k_{\mathrm{on}}[\mathrm{B}]_{\mathrm{f}}[\mathrm{W}]_{\mathrm{f}} - k_{\mathrm{off}}^{\mathrm{W:B,mis}}[\mathrm{B:W}]_{\mathrm{mis}},\\
\frac{\mathrm{d}[\mathrm{B:W}]_{\mathrm{spc}}}{\mathrm{d}t} &= Nk_{\mathrm{on}}[\mathrm{B}]_{\mathrm{f}}[\mathrm{W}]_{\mathrm{f}} - k_{\mathrm{off}}^{\mathrm{B:W,spc}}[\mathrm{B:W}]_{\mathrm{spc}},
\end{aligned}
\tag{S30}
$$

with

$$
\begin{aligned}
[\mathrm{P}] &= [\mathrm{P}]_{\mathrm{f}} + [\mathrm{P:R}] + [\mathrm{P:W}] + [\overline{\mathrm{R}}:\mathrm{R}] + [\overline{\mathrm{W}}:\mathrm{W}],\\
[\mathrm{R}] &= [\mathrm{R}]_{\mathrm{f}} + [\mathrm{P:R}] + [\mathrm{B:R}] + [\overline{\mathrm{R}}:\mathrm{R}],\\
N[\mathrm{W}] &= N[\mathrm{W}]_{\mathrm{f}} + [\mathrm{P:W}] + [\mathrm{B:W}] + [\overline{\mathrm{W}}:\mathrm{W}],\\
N[\mathrm{B}] &= N[\mathrm{B}]_{\mathrm{f}} + [\mathrm{B:R}] + [\mathrm{B:W}]_{\mathrm{spc}} + [\mathrm{B:W}]_{\mathrm{mis}}.
\end{aligned}
\tag{S31}
$$

Here, we used the following notations; $[\mathrm{P:W}] = \sum_i [\mathrm{P:W}_i]$, $[\overline{\mathrm{W}}:\mathrm{W}] = \sum_i [\overline{\mathrm{W}}_i:\mathrm{W}_i]$, $[\mathrm{B:R}] = \sum_j [\mathrm{B}_j:\mathrm{R}]$, $[\mathrm{B:W}]_{\mathrm{mis}} = \sum_{i\neq j}[\mathrm{B}_j:\mathrm{W}_i]$, and $[\mathrm{B:W}]_{\mathrm{spc}} = \sum_{i=j}[\mathrm{B}_j:\mathrm{W}_i]$. We used the parameters obtained by the melting curve analysis (Table. S4) and global fitting of the kinetic analysis (Section S5).

The values $\alpha_{\mathrm{R}}$, $\alpha_{\mathrm{W}}$ and $\eta$ are defined as $\alpha_{\mathrm{R}} = r/[\mathrm{R}]$, $\alpha_{\mathrm{W}} = \sum_i w_i/[\mathrm{W}_i]$, and $\eta = \alpha_{\mathrm{W}}/(\alpha_{\mathrm{R}} + \alpha_{\mathrm{W}})$. Here, $r$ and $w_i$ are the amount of produced copies of R and $\mathrm{W}_i$ per cycle.

### S5 Fitting of experimental data

We fitted the experimental results with various polymerase concentrations by (S13). We performed a global fitting over the three blocker conditions with a cost function

$$C(\{\mu\},\{\sigma\}|p) = \sum_{m=+\mathrm{B_R},-\mathrm{B},+\mathrm{B_W}} \sum_{l} \left[ \left( \frac{\mu^r_{l,m} - f^r_{l,m}(p)}{\sigma^r_{l,m}} \right)^2 + \left( \frac{\mu^w_{l,m} - f^w_{l,m}(p)}{\sigma^w_{m,l}} \right)^2 \right]. \tag{S32}$$

We collectively denote the fitting parameters $k_{\mathrm{on}}$, $k^{\mathrm{R}}_{\mathrm{pol}}$, and $k^{\mathrm{W}}_{\mathrm{pol}}$ as $p$. Here, $m$ and $l$ are the labels for the blocker used and polymerase concentration, respectively. The summation over $l$ is taken over all the polymerase concentrations. $\mu^{r/w}_{l,m}$ and $\sigma^{r/w}_{l,m}$ are the averages and standard deviations, respectively, of $r$ or $w$ over three replicates at each $m$ and $l$. $f^{r/w}_{l,m}$ are the values calculated by Eq. (S13).

We obtained $k_{\mathrm{on}} = 2.33 \times 10^6\,\mathrm{M^{-1}s^{-1}}$, $k^{\mathrm{R}}_{\mathrm{pol}} = 2.38 \times 10^{-2}\,\mathrm{ml/units}$, and $k^{\mathrm{W}}_{\mathrm{pol}} = 1.94 \times 10^{-3}\,\mathrm{ml/units}$ as the fitting parameters at 60 °C(Table. S5). The values of $K_{\mathrm{d}}$ were obtained by the melting analysis (Table. S4). We use $t = 30\,\mathrm{s}$, which is the annealing time in our experiments.

We discuss the validity of the obtained values of the fitting parameters. For simplicity, our model ignores details of the dynamics such as the thermal cycling and polymerase dissociation from the primer-template complex. Despite such simplification, the values of the fitting parameters are compatible with independent experimental measures from the literature. Values of $k_{\mathrm{on}}$ are typically in the order of $1 \times 10^6\,\mathrm{M^{-1}s^{-1}}$ for the DNA strands but significantly depend on the experimental conditions[4–6]. The value $k_{\mathrm{on}} = 2.33 \times 10^6\,\mathrm{M^{-1}s^{-1}}$ obtained here is of the same order of magnitude.

It is difficult to analyze the polymerization rate because we do not know the exact concentration of the active polymerase molecules. Therefore, we focus on the ratio of the polymerization rates $k^{\mathrm{R}}_{\mathrm{pol}}/k^{\mathrm{W}}_{\mathrm{pol}} = 12.3$. We are not aware of a reference value for polymerization with a mismatch in the middle of the primer. In the case of the mismatch at the 3' end of the primer, the ratio $k^{\mathrm{R}}_{\mathrm{pol}}/k^{\mathrm{W}}_{\mathrm{pol}}$ was estimated to be at least larger than 50[7]. It is reasonable that the present value is lower than that value because the mismatch at the 3' end is known to cause a more severe hindrance to the polymerase reaction than the internal mismatch.

We also did the same experiment at 57.5 °C and fitted the results with our model to obtain parameters at this temperature. We obtained $k_{\mathrm{on}} = 5.07 \times 10^6\,\mathrm{M^{-1}s^{-1}}$, $k^{\mathrm{R}}_{\mathrm{pol}} = 2.98 \times 10^{-2}\,\mathrm{ml/units}$, and $k^{\mathrm{W}}_{\mathrm{pol}} = 2.26 \times 10^{-3}\,\mathrm{ml/units}$ as the fitting parameters (Fig. S6). $K_{\mathrm{d}}$ at 57.5 °C are shown in the Table. S4. The annealing time $t$ was 30 s. The value of $k_{\mathrm{on}}$ was similar to that of 60 °C. The values of $k^{\mathrm{R}}_{\mathrm{pol}}$ and $k^{\mathrm{W}}_{\mathrm{pol}}$ were larger than that of 60 °C, which could be explained as the stabilization between the primer and the templates.

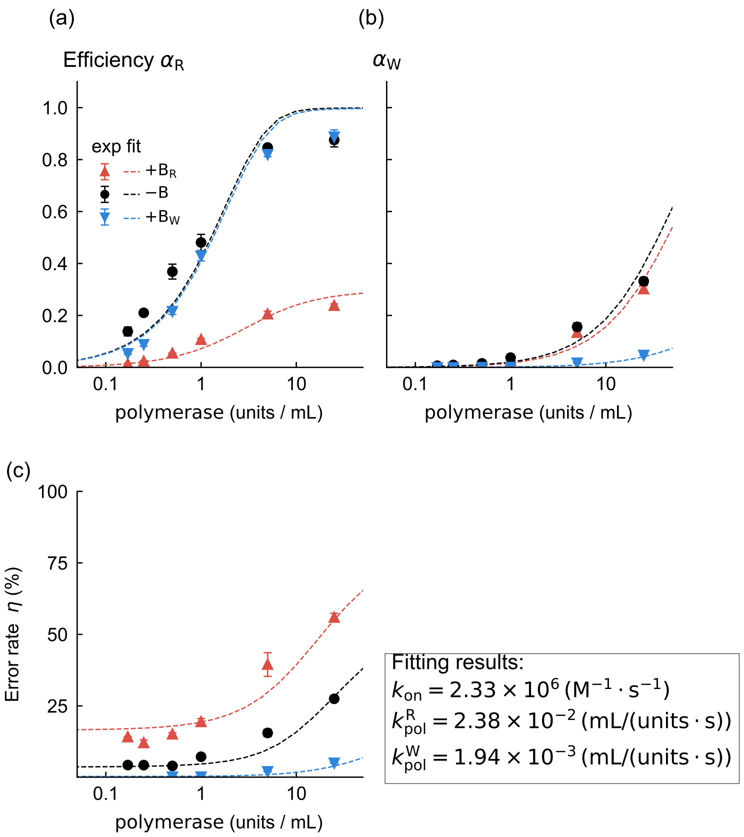


**Figure S5.** Results of the kinetic analysis at 60 °C in linear scale. Best fit parameters are shown in the box. The error bars represent standard deviations.

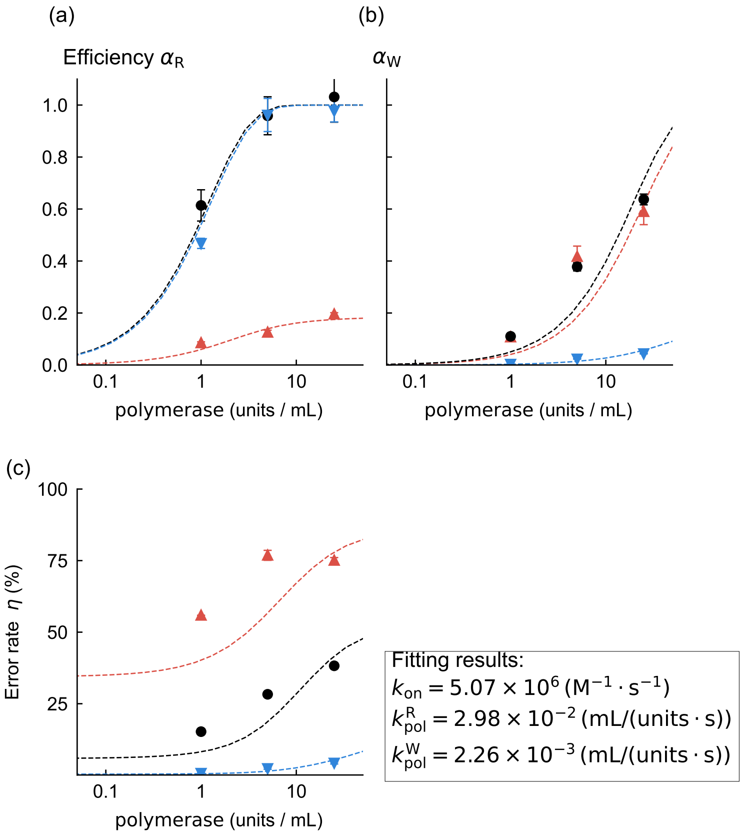


**Figure S6.** Scatters are experimental results of the kinetic analysis at 57.5 °C. The error bars represent standard deviations ($n = 3$). The polymerase concentration we tested were 1, 5, and 25 units/mL. The annealing time was 30 s. The number of cycles was ten at all the polymerase conditions. Dashed lines are the global fitting results. Best fit parameters are shown in the box.

### Supplementary Figures

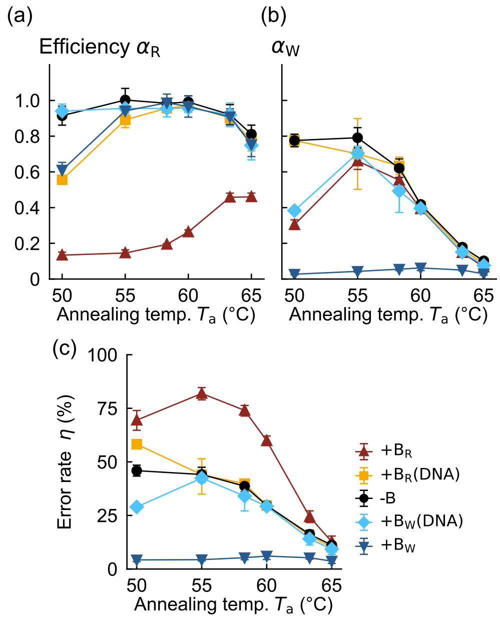


**Figure S7.** The results of changing the annealing temperature $T_a$ with the LNA-free blocker. The error bars represent standard deviations ($n = 3$).
